# Particle fusion of Single Molecule Localization Microscopy data reveals dimer structure of Nup96 in Nuclear Pore Complex

**DOI:** 10.1101/2022.10.04.510818

**Authors:** Wenxiu Wang, Arjen Jakobi, Yu-Le Wu, Jonas Ries, Sjoerd Stallinga, Bernd Rieger

## Abstract

Single molecule localization microscopy offers nowadays resolution nearly down to the molecular level with specific molecular labelling, thereby being a promising tool for structural biology. In practice, however, the actual value to this field is limited primarily by incomplete fluorescent labeling of the structure. This missing information can be completed by merging information from many structurally identical particles equivalent to cryo-EM single-particle analysis. In this analysis, we present particle averaging of fluorescently labelled Nup96 in the Nuclear Pore Complex followed by data analysis to show that Nup96 occurs as a dimer with in total 32 copies per pore. We use Artificial Intelligence assisted modeling in Alphafold to extend the existing cryo-EM model of Nup96 to accurately pinpoint the positions of the fluorescent labels and show the accuracy of the match between fluorescent and cryo-EM data to be better than 3 nm in-plane and 5 nm out-of-plane.

## 1. Introduction

The Nuclear Pore Complex (NPC) is an essential molecular machine embedded in the nuclear envelope connecting the nucleus to the cytoplasm (D’Angelo and Hetzer, 2008). The NPC is indispensable in eukaryotic cellular processes, such as regulating the transportation of protein and ribonucleoprotein (Kabachinski and Schwartz, 2015; Wente and Rout, 2010), generating a diffusion barrier that separates the nuclear compartment from the cytoplasm (Hoelz et al., 2011; Knockenhauer and Schwartz, 2016) and working as gateway for gene regulation (D’Angelo, 2018). The structure and molecular composition of the NPC, in particular the scaffold, has been extensively studied. Previous cryogenic electron microscopy (cryo-EM) studies have resolved the structure of the NPC scaffold to high-resolution. The scaffold is composed of multiple copies of about 34 different nucleoporins (Nups) and most of the Nups are organized in two rings each showing eight-fold rotational symmetry (Hoelz et al., 2016; Lin et al., 2016; Mi et al., 2015; Cronshaw et al., 2002; Rout et al., 2000). In the cryo-EM map of the human NPC scaffold of von Appen et al. (Appen et al., 2015) each ring contains 16 Nup96 molecules organized into dimers with eight-fold symmetry.

Super-resolution microscopy is emerging as a complementary technique to study biological structure since it enables ‘diffraction unlimited’ resolution (Hell, 2009; Klein et al., 2014; Vicidomini et al., 2018). Single molecule localization microscopy (SMLM) is one of these super-resolution techniques and obtains super-resolved images with a resolution of 20 nm by localizing single fluorescent emitters (Lelek et al., 2021; Hell, 2009). If many chemically identical structures, called particles, can be imaged, they can be registered and combined into one “super-particle”. With this strategy, the often poor degree of labelling of each individual particle can be mitigated and an even better resolution can be obtained (Löschberger et al., 2012; Szymborska et al., 2013; Broeken et al., 2015; Gray et al., 2016). Different 2D template-based particle fusion methods have been applied in SMLM to demonstrate the eight-fold rotational symmetry of the NPC (Löschberger et al., 2012, 2014). These methods, however, carry the risk to generate reconstructions with a bias toward the template. Later, a template-free 2D registration approach could reveal the eight-fold symmetry of the NPC in an unbiased manner (Heydarian et al., 2018). This template-free method was extended to 3D and used to reconstruct the 3D structure of Nup107 and Nup96 revealing 2 phase shifted rings with eight blobs per ring (Heydarian et al., 2021). Yet, the 3D approach by Heydayrian et al. suffered from the ‘hot spot’ artefact, which could only be mitigated by applying prior knowledge about the eight-fold symmetry in a post-processing step. In another template-free 3D particle fusion approach, the super-particle is generated based on a data-driven template derived by pairwise similarities of individual particles (Wu et al., 2021). This approach also shows two rings with each 8 blobs or clusters in the NPC reconstruction, as expected, but interestingly some of the blobs are elongated and tilted in the plane of the rings. Limited by their high computational cost, neither the approach by Heydarian et al. (2021) nor by Wu et al. (2021) can reconstruct thousands of particles in a reasonable time. We recently introduced a template free and fast particle fusion approach (Wang et al., 2022) that overcomes the ‘hot spot’ problem and the limitation to computation speed, so that datasets of several thousands of particles (or more) are now accessible for structural analysis.

Up to now SMLM of the NPC was able to reveal the eighth-fold symmetry, resolve 8 spots individually per ring and could show the separation of the nuclear and cytoplasmic rings in 3D. Although from cryo-EM work (Appen et al., 2015) it is known that e.g. Nup96 should occur as a dimer, this could yet not be resolved by SMLM. Here, for the first time, we show that each of the 8 blobs indeed contains two fluorophores attached via a SNAP tag to the Nup96 dimer by combining our fast template free particle averaging method (Wang et al., 2022) with careful data analysis on 5 datasets (originating from five nuclei of five cells) of in total 4,538 NPCs (Wu et al., 2021). We made a detailed comparison of the outcome of our analysis to the cryo-EM data. To this end, we extended the incomplete Nup96 model derived from von Appen et al. (Appen et al., 2015) by Alphafold (Jumper et al., 2021) to find the positions where the fluorescent SNAP-tags are expected to attach to the Nup96. Next, we registered our estimated positions of the fluorescent dimers to these expected positions of the SNAP-tag from the cryo-EM model and found the average distance between the SNAP positions derived from the cryo-EM model and from our SMLM data analysis to be < 3 nm laterally and 5 nm axially.

## 2. Methods

### 2.1. Particle Averaging

We applied our previously published particle averaging method (Wang et al., 2022) to five NPC SMLM datasets individually and all 5 combined (total of 4,538 NPCs). Dataset 1 with 368 NPCs was previously described (Wu et al., 2021) and the other datasets were obtained in the same way. We use the default values given in Wang et al. (2022) for the data fusion algorithm, except for the number of Gaussian components *K* in the Gaussian Mixture Model (GMM) and the initial Gaussian standard deviation of the Gaussian components. We set *K* = 34, that is 32 Gaussian components for 32 binding sites and 2 to accommodate false positive localizations of the nuclear ring (NR) and the cytoplasmic ring (CR). The initial poses of the NPC particles are roughly aligned with the optical axis, as the nuclear membrane runs along the cover slip. We can use this to set an initial Gaussian standard deviation to 33 nm, smaller than the overall size of the NPC, to make the algorithm converge faster. We obtained super-particles with two rings each showing 8 elliptically shaped blobs for all 5 datasets, as well as for the combined dataset (Fig. 1a-c).

**Figure 1:**
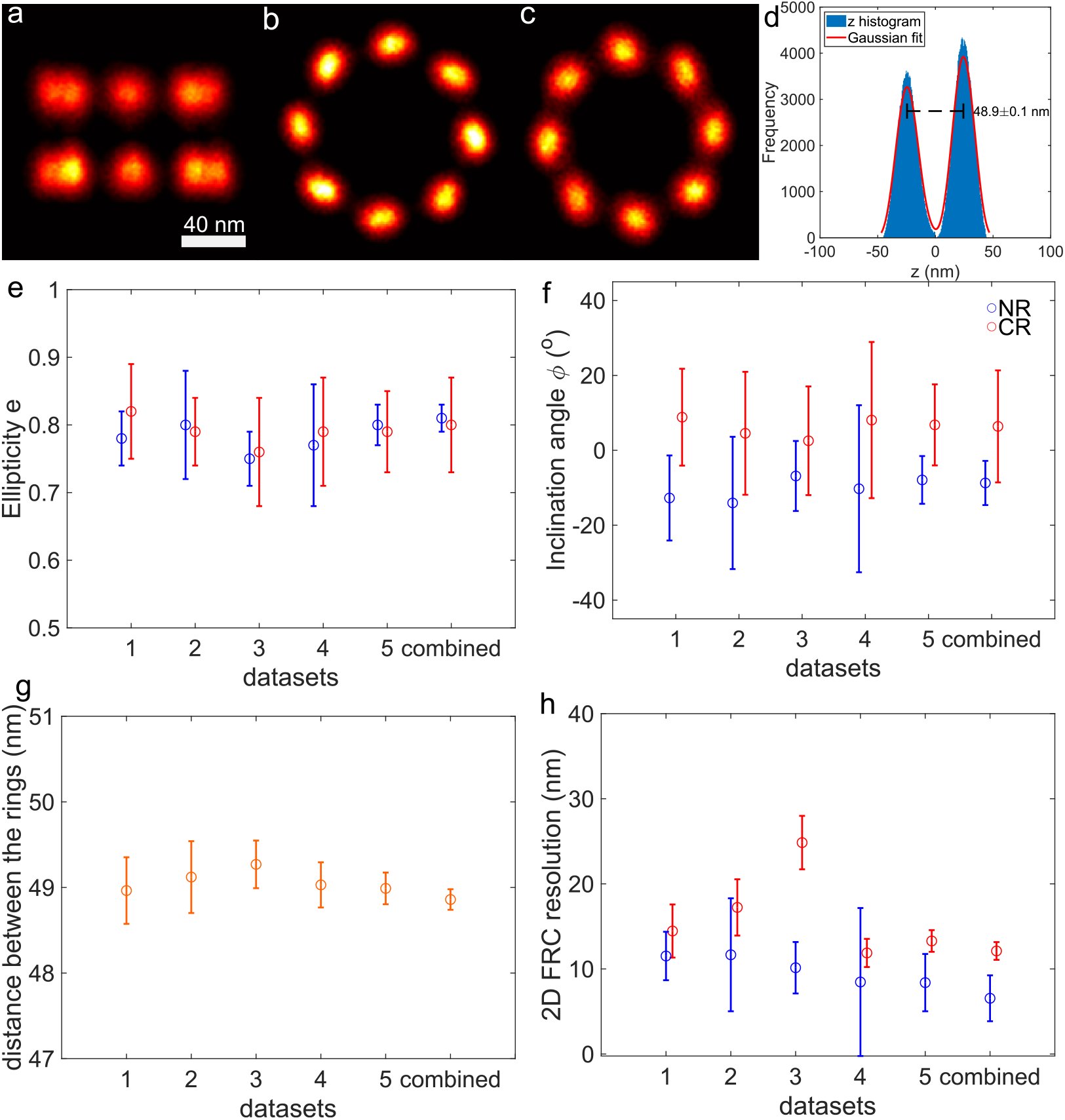
(a-c) 3D reconstruction of Nup96 from 4,538 NPC particles of the combined dataset. The reconstruction resolves two rings and 8 blobs per ring. (a) Side view, (b) top view of the nuclear ring (NR), (c) top view of the cytoplasmic ring (CR), (d) histogram of *z* coordinates of localizations in the reconstruction obtained from the combined dataset, and bimodal Gaussian fit to the data. The distance between the two peaks of the histogram is 48.9 ± 0.1 nm (e) Average ellipticity *e* of the elliptical blobs for the NR (blue) and CR (red), (f) Average angle *ϕ* of the long axis of the elliptical blobs with the tangent to the ring in the *xy*-plane for the NR (blue) and CR (red) (negative angles indicate clockwise rotation). (g) Distance between the NR and CR for all the datasets. (h) 2D FRC resolution of the NR and CR for all the datasets. Scale bar in (a) is 40 nm and applies to (b,c) as well. All figures present results after removing outliers.

### 2.2. Splitting in two rings

We obtain super-particles from the particle averaging of the different datasets that have the same shape but different absolute poses in 3D space. For better comparison and easier analysis we align their global pose such that rings are perpendicular to the *z*-axis of our global coordinate system. We do so by registering the super-particles to a fixed template which has 2 rings and each ring contains 8 points with 8-fold symmetry. The center of the template locates at the origin of the coordinate system. The distance between the two rings in the template is 50 nm and the radius of each ring is 55 nm. Finally, we rotate the super-particles around the *x*− and *y*−axis from −2^*o*^ to −2^*o*^ in steps of 0.1^*o*^ to find the minimal full width at half maximum (FWMH) of the histogram of *z*-coordinates. Then, we split the super-particle into two 8 blob rings according to their *z*-coordinates for further analysis. These rings represent the nuclear (NR) and cytoplasmic (CR) ring of the NPC. The reconstruction quality is assessed by computation of the 2D FRC resolution (Nieuwenhuizen et al., 2013) separately for the NR and CR.

### 2.3. Outlier localization filtering

Next, we remove outlier localizations in each ring. To this end we fit the particle to a randomly initialized GMM with 9 Gaussian components using the Joint Registration of Multiple Point Clouds (JRMPC) method (Evangelidis and Horaud, 2017). The Gaussian components obtained from this fit have different standard deviations and we find outlier localizations by comparing the standard deviations of the components. In all cases we find that there are 8 components with similar (small) standard deviation and one component with a large standard deviation, indicating the outlier component. We remove the localizations in the latter component from the data. As the automated data analysis is sensitive to outlier localizations, we apply further outlier filtering based on the density of localizations to the rings. We remove all localizations with a local density smaller than half of the mean density value of a single ring such that every blob is separated clearly from its adjacent blobs in the reconstruction.

### 2.4. Ellipticity measurement in per blob

The long axes of the ellipsoidally shaped blobs in a ring are tilted with respect to the circumference of the ring. To investigate this further, we measure the ellipticity of each projected blob onto the *xy*-plane. The ellipticity is defined as *e* = *b/a*, where *a* and *b* are the length of the long and short axis, respectively. We also measure the inclination angle between the long axis of each blob and the tangent to the overall 2D ring. We then calculate the average value and standard deviation of the ellipticity and inclination angle over the set of values for the 8 blobs in each ring.

### 2.5. Anisotropic Gaussian mixture fitting to 8 blobs per ring

The inclination of the elliptical shape of the blobs with respect to the rings indicates that more than one emitter is present per blob. As moreover the inclination angle is also opposite in sign for the NR and CR, the inclination is not a reconstruction artifact. The dimer, however, is not resolvable due to the limited localization precision, residual drift or registration error. The imperfection of the pore with corners slightly moved, the distortion of the entire pore, and biological heterogeneity could also affect the resolution of the reconstructions. We assume that there are two SNAP emitters per blob based on the prior knowledge from the cryo-EM model that there are two Nup96 per blob. We again use a GMM with 16 components to fit the 8 blobs per ring and interpret the Gaussian components’ centers as the positions of the emitters. For the fitting procedure we use the following heuristics: The initial standard deviation for the Gaussian fitting is equal in *x* and *y* direction (in-plane), but two times larger in the *z*-direction (which is aligned with the optical axis), because the axial localization uncertainty is typically two to three times larger than in the *xy*-plane (Rieger and Stallinga, 2014). Furthermore, we assume that all Nup96 dimers in a ring are identical, i.e. the Gaussian components in the GMM have identical diagonal covariance matrices.

We use iterative Expectation-Maximization (EM) to find the optimal GMM by maximizing the likelihood for the GMM to fit the localizations (McLachlan and Peel, 2000). The shared diagonal covariance matrices, centers of Gaussian components and posterior probabilities of component memberships are the fitting parameters and updated iteratively. The obtained Gaussian centers from fitting 16 anisotropic Gaussian mixture model are represented by 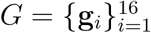 with **g**_*i*_ = (*g*_*ix*_, *g*_*iy*_, *g*_*iz*_).

### 2.6. Incorporation of the eight-fold symmetry into the fitting

We have employed a second fitting method to make use of the eight-fold symmetry, as unconstrained anisotropic GMM fitting is sensitive to the setting of the initial Gaussian centers. To this end we generate 16 points with eight-fold symmetry as the initial centers of the anisotropic GMM fitting for every ring. The 16 binding sites defined by the point set 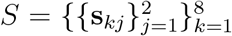 are characterized by 6 parameters: (*c*_1*x*_, *c*_1*y*_, *c*_1*z*_, *d*, *θ*, *ϕ*), where **c**_1_ = (*c*_1*x*_, *c*_1*y*_, *c*_1*z*_) is the center of the first (*k* = 1) dimer, *d* is the distance between the two binding sites in the dimer, *θ* is the angle between the line connecting the two binding sites in a dimer and the positive *z*-axis (0 ≤ *θ* ≤ *π*), and *ϕ* is the angle between the projection of this line on the *xy*-plane to the tangent of the ring in *xy*-plane (0 ≤ *ϕ* ≤ 2*π*). The in-plane coordinates of the dimer center can be parameterized as **c**_1_ = (*c*_1*x*_, *c*_1*y*_) = *R* (cos *ψ*, sin *ψ*), with *R* the radius of the ring and *ψ* the in-plane angle (compare Fig. Appendix A). The coordinates of the two binding sites in the first dimer are thus given by:

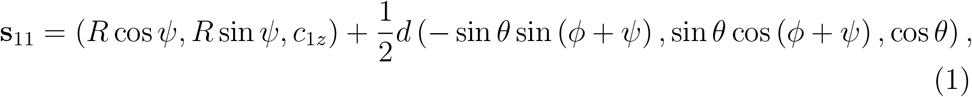

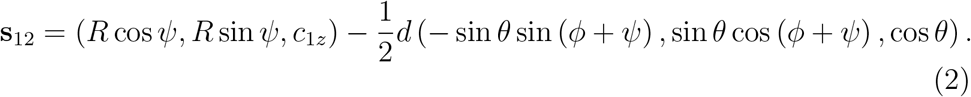

We choose emitter 1 to be the emitter that lies above the center plane of the ring, i.e. we restrict the polar angle to 0 ≤ *θ* ≤ *π/*2. The coordinates of the binding sites of the *k* = 2, …, 8 other dimers can be derived by rotating these points by (*k* − 1)*π/*4, *k* = 2, … 8 in the *xy*-plane. The parameters for the initial GMM centers are (0, 53.5 nm, ±25 nm, 13 nm, *π/*2, *ϕ*), where + corresponds to NR and − to CR. The in-plane angle *ϕ* is randomly generated in the interval 0 to *π*. We find the parameters of the symmetry constrained point sets *S* from the coordinates of the unconstrained point set *G* by minimizing the mean square error between the point sets *S* and *G* with the quasi-Newton method (Shanno, 1970) implemented in MATLAB. We repeat the processing of anisotropic Gaussian mixture fitting and incorporation of the eight-fold symmetry 100 times with different initial centers to obtain an uncertainty measure for the fits

### 2.7. Comparison with the cryo-EM model

In order to relate the found emitter positions to available structural data on the human NPC (Appen et al., 2015), we modeled the full-length Nup96 structure using Alphafold2 (Jumper et al., 2021) and added the SNAP-tag at the carboxyl-terminal position on the Nup96s. We rigid-body fitted the full-length Nup96 into the cryo-EM density using residues 881-1817 from PDB ID 5a9q as anchor residues (Appen et al., 2015). As expected, the SNAP tag protrudes from the NPC assembly, adding support for its correct placement. We then symmetry-expanded the fitted model to generate the full NPC assembly and denote the center-of-mass of the O^6^-benzylguanine-AF647 (BG-AF647) to represent the emitter position.

The fitted eight-fold symmetric emitter positions of the combined dataset are registered with the SNAP tags from the cryo-EM model by JRMPC. The predicted SNAP tag positions based on the cryo-EM model are represented by position vectors **m**_*kj*_ = (*m*_*kj,x*_, *m*_*kj,y*_, *m*_*kj,z*_) for *k* = 1, … 8 and *j* = 1, 2. Using this set of position vectors reference values for the radius *R* of the CR and NR, the distance *d* between the two binding sites in the dimer, the in-plane inclination angle *ϕ* of the dimer, and the out-of-plane tilt angle *θ* of the dimer can be obtained from equations similar to Eqs. (1) and (2).

The two models can be directly compared by computing the in-plane and axial distance of the cryo-EM SNAP tag positions to the estimated emitter positions from SMLM particle fusion:

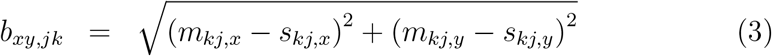

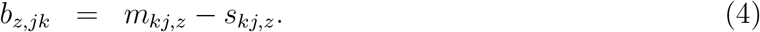

As the cryo-EM data satisfies the eight-fold rotational symmetry by construction we only compare the cryo-EM SNAP tag positions to the estimated emitter positions from SMLM particle fusion that are obtained from the symmetry constrained dimer fits. That means that there are only four distinct values for *b*_*xy,jk*_ and four distinct values for *b*_*z,jk*_ (CR and NR, two emitter positions per dimer).

## 3. Results

We have applied the above described methodological steps to the five Nup96 datasets. Dataset 1 was described earlier by (Wu et al., 2021); datasets 2-5 result from the same cell line and staining and imaging protocol(Thevasthasan et al., 2019). The number of analyzed NPCs per dataset are 368, 568, 706, 1178 and 1,718 for a total of 4,538 NPCs.

### 3.1. Particle fusion indicates inclined elliptical blobs

In Fig. 1a-c we show the super-particle reconstruction of the combined dataset consisting of 4,538 NPCs. In the top-view of the nuclear (NR) and cytoplasmic (CR) ring (b,c) the elongation of the blobs is clearly visible. The average ellipticity of the blobs is about 0.8 over all datasets (Fig. 1e). Furthermore, it appears that the long axis of the ellipses makes an angle of around ±8° with the tangent to the rings, with opposite sign for the CR and NR (Fig. 1e). If the ellipticity was due to a registration error or artefact of the particle fusion procedure, then the long axis of the ellipses would be aligned to the tangent of the rings. We can therefore conclude that the non-zero inclination of the elliptical blobs is evidence of a structural feature. The most simple explanation is that each elliptical blob is the composite of localization events of emitters that bind to two distinct binding sites, i.e. the particle fusion directly suggests that Nup96 occurs in a dimer arrangement.

We note that the evidence for (at least) two binding sites comes from interpreting the structural feature of ellipticity. The binding sites cannot be directly observed as two distinct spots in the reconstructions as the 2D FRC resolution is just 9.5 ± 2.0 nm for the NRs and 15.6 ± 4.9 nm for the CRs (see Fig. 1h), which is comparable to the expected distance between the two binding sites of around 12 nm (Appen et al., 2015).

### 3.2. Unconstrained and symmetry constrained fitting of binding sites

We investigate two approaches to estimate the positions of the two binding sites in the Nup96 dimers. Firstly, we use unconstrained fitting of 16 anisotropic Gaussian centers for the NR and CR separately, and secondly we use constrained fitting in which we impose the eight-fold rotational symmetry (according to Eqs. (1) and (2)).

We find the average distance for the combined data to be 1.93 ± 0.43 nm for the NR and 1.84 ± 0.69 nm for the CR. This small difference indicates that our data matches well with the eight-fold symmetry and that the use of the symmetry constraint in the fit is a valid procedure.

### 3.3. Structural parameters of the Nup96 dimer

Fig. 3 shows the structural parameters (radius *R* of CR and NR, distance *d* between the two binding sites in the dimer, in-plane inclination angle *ϕ* of the dimer, out-of-plane tilt angle *θ* of the dimer) for all datasets for the case of symmetry constrained fitting. For comparison we also show the reference values from the cryo-EM data (*R* = 54 nm, *d* = 11.8 nm, *ϕ* = −32.6° (NR), *ϕ* = 32.1° (CR), *θ* = 76.8°). The found structural parameters are relatively consistent between the different datasets and all the parameters are close to the cryo-EM reference values. In particular the radius and distance between the binding sites in the dimer match well, where the angles are a bit more off compared to the cryo-EM model. There are some variations between the datasets, where datasets 3 and 4 appear to have more scattered localizations than the others resulting in visually poorer reconstructions and FRC values for the NR and CR. The estimated uncertainties of the parameters for the NR are smaller than for the CR, which could originate from the way JRMPC fuses the data. In a global optimization part of the structure (here NR) could be matched better at the expense of CR alignment. The radius of the NR matches very well with the cryo-EM reference value, whereas the radius of the CR is consistently about 3 nm smaller. The estimate of the distance between the binding sites in the dimer is somewhat smaller than the cryo-EM reference distance. We find that the in-plane inclination angle has the opposite sign for the NR and the CR, which is consistent with the cryo-EM model. The estimated magnitude of the in-plane inclination angle is somewhat smaller than the cryo-EM reference values. A larger quantitative mismatch is found for the out-of-plane tilt angle, which we attribute to the localization uncertainty in the axial direction, which is two to three times larger than the uncertainty in the *xy*-plane (Rieger and Stallinga, 2014).

**Figure 2:**
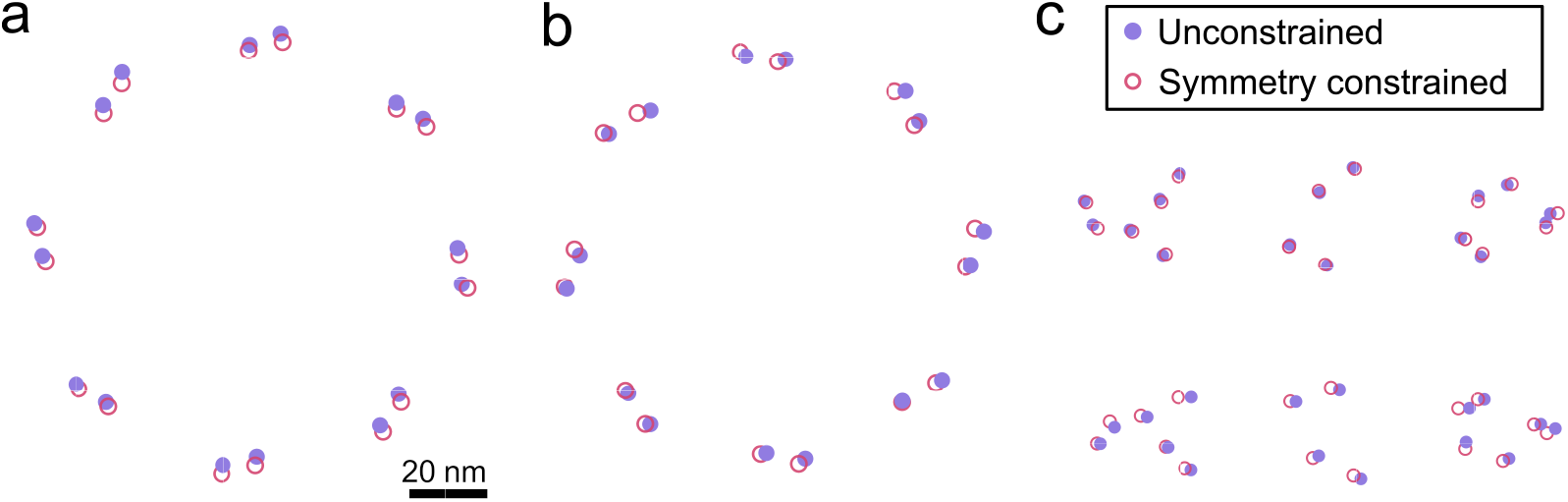
Locations of binding sites obtained from an unconstrained fit (purple) and from an eight-fold symmetry constrained fit (pink) of the combined data. We fit 16 sites for the nuclear (NR) and cytoplasmic ring (CR) separately. Top view of binding sites for the NR (a) and CR (b). (c) Oblique view of CR and NR for projection at an angle 15^*o*^ with the *xy*-plane. Scale bar in (a) applies to (b,c) as well.

**Figure 3:**
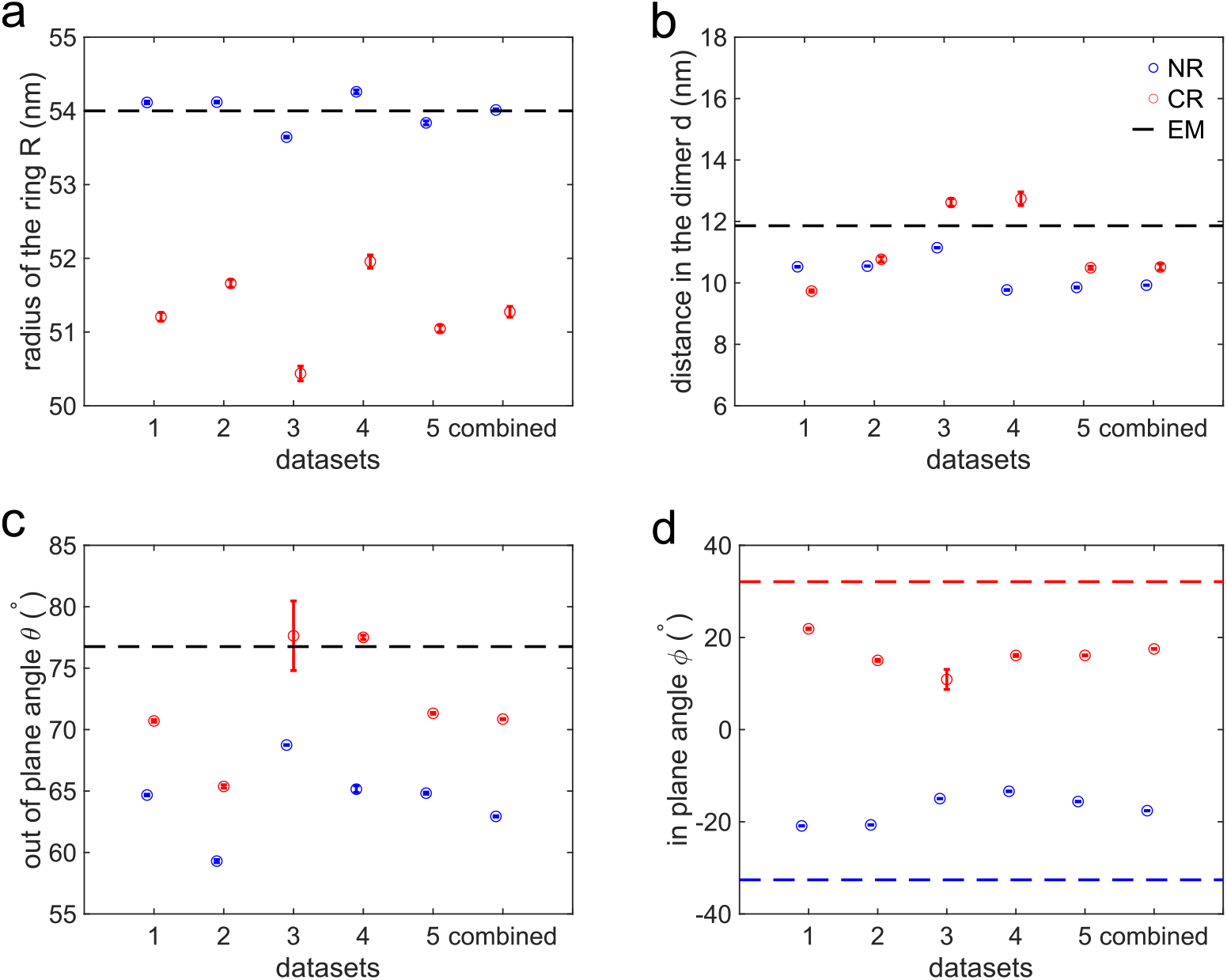
Structural parameters for eight-fold symmetry constrained fitting for the NR (blue) and CR (red) for all datasets. Parameters of fluorophore locations derived from the cryo-EM model are shown as dashed lines. (a) Radius for the NR and CR, the NR radius is around 54.0 nm and the CR radius is around 51.2 nm. The reference radius is 54.2 nm. (b) Distance between the binding sites in the dimers. The distance in the dimers is 10.7 ± 1.0 nm and the reference value is 11.8 nm. (c) Out-of-plane angle *θ* of the line connecting the two emitters in a dimer to the *z*-axis, *θ* = 68.3° ± 5.6° and the reference *θ* = 76.8°. (d) In-plane angle *ϕ* of the line connecting the two emitters in the dimer to the tangent of the ring in the *xy*-plane, *ϕ* ∼ −17.2° for the NR and *ϕ* ∼ +16.3° for the CR. The reference angle is −32.6° for the NR (blue dashed line) and 32.1° for the CR (red dashed line).

### 3.4. Comparison of cryo-EM and SMLM positions

Figure 4 shows a direct comparison between the positions of the SNAP tags predicted from the cryo-EM data, and the positions of the emitters according to the SMLM particle fusion data. For the NR we find in-plane distances of 3.0 nm and 1.4 nm, and out-of-plane distance of 2.8 nm and 5.6 nm for the two emitter positions in the dimer, for the CR we find in-plane distances of 2.3 nm and 3.6 nm, and out-of-plane distance of −5.4 nm and −4.5 nm for the two emitter positions in the dimer. Please note that we cannot assign the fluorophore position directly in our structural model, which is limited to modeling the SNAP tag position relative to Nup96, whereas we measure the fluorophore position directly in SMLM. The expected distance of the SNAP-tag and the fluorophore position is 1-2 nm. Overall the agreement in the plane of the NPC rings is rather well, with an overall error of just 2.6 ± 0.9 nm.

**Figure 4:**
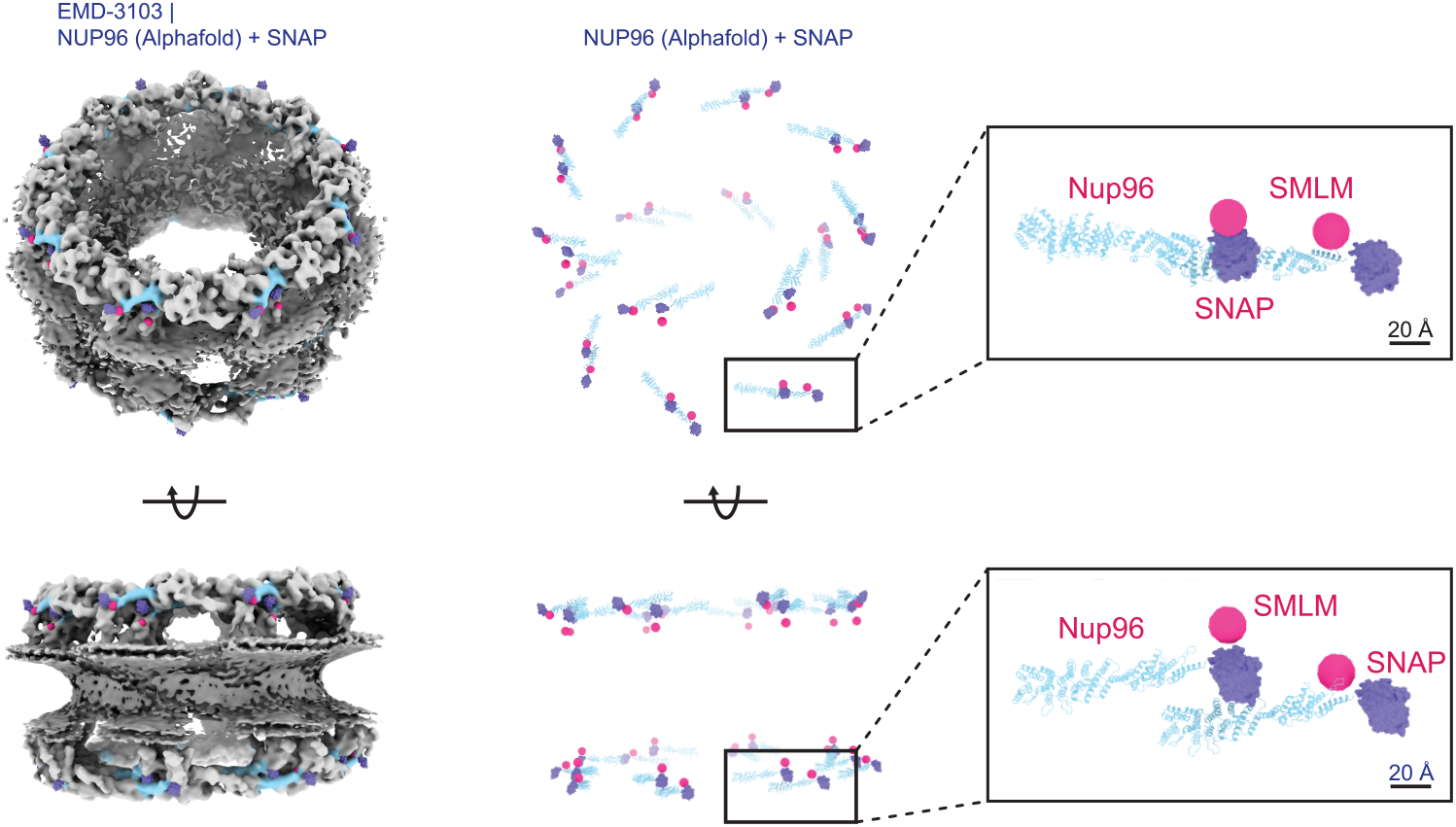
Overlay of the fluorophore positions from the SMLM particle fusion data (pink) and the SNAP-tag derived from the cryo-EM data (purple).

The lateral error also seems free of a systematic bias between the cryo-EM based and the SMLM particle fusion based position estimates. This stands in contrast to the axial position estimates where we find a distance between the top emitters in the CR and the NR of 8.2 nm, and between the bottom emitters in the CR and NR of 10.1 nm. It appears that the SMLM data give a distance between the NR and CR that is systematically smaller than the distance obtained from the cryo-EM model, with an average bias in estimated NPC thickness obtained from the cryo-EM/SMLM comparison of 9.2 nm. Fig. 1g) shows the estimated distance between the NR and CR for all datasets, as well as the histogram of the *z* coordinates of the 275,809 localizations in the reconstruction obtained from the combined dataset. We find the distance between the NR and CR in SMLM to be 48.9 ± 0.1 nm, while the distance between the rings in the EM model is 57.2 ± 0.2 nm, a bias of about 8.3 nm. This analysis of the underlying axial localization data shows that the bias found is not due to an artefact of the particle fusion method.

## 4. Discussion

Overall, our analysis of the SMLM particle fusion based data points to a Nup96 dimer structure that matches well with the cryo-EM data. Compared to the cryo-EM model, however, one major inconsistency remains. Namely, the height of the NPC is estimated from SMLM particle fusion to be about 8 nm less than from cryo-EM. Similar ring separation values for Nup96 have been reported in other SMLM studies (Gwosch et al., 2020; Thevasthasan et al., 2019), and have a larger deviation from the cryo-EM model than the levels of statistical error. There are a number of factors that could contribute to this discrepancy. Firstly, the missing wedge problem of cryo-EM reconstructions, in combination with the dominant axial orientation of the NPCs in cryo-EM imaging, may compromise axial distance estimation from the cryo-EM reconstructions. Secondly, differences in sample preparation for electron and light microscopy may play a role. Anisotropic stiffness of the NPC in combination with the lower density of vitreous ice (∼ 0.94 g/cm^3^) compared to liquid water, may result in anisotropic expansion of the NPC in cryo-EM imaging. Thirdly, the axial localization data is calibrated based on reference images of NPCs that are oriented sideways, and therefore show two rings clearly separated in the image plane (Thevasthasan et al., 2019). If the fluorophore motion is restricted during imaging, dipole orientation effects on single molecule imaging can have an impact on the lateral position estimation, resulting in biases on the order of ∼10 nm (Stallinga and Rieger, 2012). A confounding factor is the analysis of 3D SMLM data of Nup107, which is adjacent to Nup96 in the NPC scaffold, and which was found to have a ring separation of 60 nm in SMLM particle fusion (Heydarian et al., 2021), in agreement with the cryo-EM data.

The estimated structural parameters for the NR appear to be more consistent between datasets than for the CR (see Fig. 3). We attribute this to the fact that the localizations of the CR are more scattered, and in turn the reconstructed super-particle is of lower quality there (FRC resolution equal to 10.8 ± 1.5 nm for the NRs and to 17.0 ± 5.5 nm for the CRs). The underlying reason could be that the 3D particle fusion favors alignment of one ring, which leaves the other ring more blurred - in particular in the axial direction. A possible point of improvement could be to re-register the localizations in the CR by our particle fusion method separately, which in the end may give rise to a reconstruction of the CR with better quality.

Other technical improvements to the data analysis can also be envisioned. Firstly, the removal of outlier localizations is now done via a cascade of adding an extra Gaussian center to the GMM and subsequently filtering localizations according to the density of localizations. A single integrated approach may improve robustness of the data analysis procedure. Secondly, initial parameter settings could result in a better convergence to global optima of GMM fitting. A more principal refinement of the current analysis relates to model selection. The current analysis relies on the simplest model that can explain the observations, namely that the Nup96 appears in a dimer structure. A possible improvement may therefore be found in a statistical criterion that supports that there are 32 emitters in the NPC, as opposed to another multiple of 16.

Recently, Helmerich et al. (2022) speculated that fluorophores at distances below about 10 nm cannot be reliably resolved with SMLM although the precision given by the microscope, data analysis pipeline, and observed single molecule photon count should allow such a distinction. Energy transfer between the close-by fluorophores is assumed to cause re-excitation of the emitters in the beginning of the experiment followed by bleaching of both, preventing observation of them individually. In particular with the introduction of MINFLUX and related techniques (Balzarotti et al., 2016; Gwosch et al., 2020; Cnossen et al., 2020; Jouchet et al., 2021) that offer nanometer localization precision at low photon count these observations could frustrate direct imaging of the dimer separation in the NPC except for rare cases and leave careful data analysis as presented here as the only remaining tool.

In conclusion, we have demonstrated that single molecule light microscopy can reveal 3D structures in the small nanometer range. Our high-resolution particle fusion reconstructions and subsequent data analysis enabled the precise estimation of the positions of 32 emitter sites for Nup96 in the NPC. The comparison with cryo-EM data shows consistency better than 3 nm laterally, but also a bias in the NPC height estimation of about 8 nm. The latter inconsistency is ill understood, and requires further study by the research community.

## 5. Acknowledgements

This work has been supported by the Dutch Research Council (NWO), VICI grant no. 17046 for B.R. and W.W.

## 6. Author Contributions

W.W. made numerical analysis and model fits. A.J. provided the analysis of the cryo-EM data. S.S. and B.R. supervised the study. J.R. and Y.W. provided Nup96-SNAP SMLM data. All authors contributed to writing the paper.

## Appendix A. Schematic for first pair of dimers of 16 points with eight-fold symmetry

**Figure A.5:**
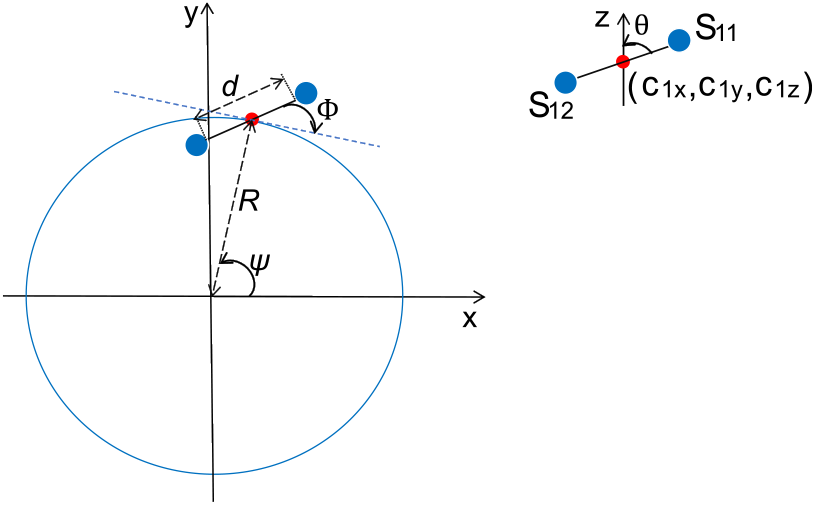
Schematic for first pair of dimers of 16 points with eight-fold symmetry. The red point is the center of the first pair of dimers and the blue points are dimers.

